# Disrupted Peyer’s patch microanatomy in COVID-19 including germinal centre atrophy independent of local virus

**DOI:** 10.1101/2021.12.17.473179

**Authors:** Silvia C. Trevelin, Suzanne Pickering, Katrina Todd, Cynthia Bishop, Michael Pitcher, Jose Garrido Mesa, Lucia Montorsi, Filomena Spada, Nedyalko Petrov, Anna Green, Manu Shankar-Hari, Stuart J.D. Neil, Jo Spencer

**Affiliations:** Peter Gorer Department of Immunology, School of Immunology and Microbial Sciences, King’s College London, London, UK; Department of Infectious Diseases, School of Immunology and Microbial Sciences, King’s College London, London, UK; NIHR Guy’s and St. Thomas Biomedical Research Centre at Guy’s and St. Thomas NHS Foundation Trust and King’s College London, London, UK; Department of Histopathology, Guy’s and St. Thomas NHS Foundation Trust and King’s College London, London, UK; Centre for Inflammation Research, Queen’s Medical Research Institute, The University of Edinburgh, Edinburgh, UK

**Keywords:** Germinal centre, Peyer’s Patch, atrophy of lymphoid follicle, gut Sars-Cov2 infection, severe COVID-19

## Abstract

Confirmed SARS-coronavirus-2 infection with gastrointestinal symptoms and changes in microbiota associated with coronavirus disease 2019 (COVID-19) severity have been previously reported, but the disease impact on the architecture and cellularity of ileal Peyer’s patches (PP) remains unknown. Here we analysed *post-mortem* tissues from throughout the gastrointestinal (GI) tract of patients who died with COVID-19. When virus was detected by PCR in the GI tract, immunohistochemistry identified virus in epithelium and lamina propria macrophages, but not in lymphoid tissues. Immunohistochemistry and imaging mass cytometry (IMC) analysis of ileal PP revealed depletion of germinal centres (GC), disruption of B cell/T cell zonation and decreased potential B and T cell interaction and lower nuclear density in COVID-19 patients. This occurred independent of the local viral levels. The changes in PP demonstrate that the ability to mount an intestinal immune response is compromised in severe COVID-19, which could contribute to observed dysbiosis.

## Introduction

Dysregulated immune response to infection with SARS-coronavirus-2 (SARS-CoV-2) is the main driver of mortality in coronavirus disease 2019 (COVID-19) (1, 2). Whilst respiratory dysfunction is common, symptoms involving the gastrointestinal (GI) tract has been identified, including vomiting and diarrhoea in 12% of the patients (3). Moreover, viral RNA has been found in stool samples (4) and viral particles identified in ileal epithelium (5). The receptor for SARS-CoV-2 angiotensin converting enzyme 2 (ACE2) is expressed on the luminal surface of epithelial cells throughout the GI tract. It has been proposed that reservoirs of virus in the GI tract could support longer lived antibody responses that are fundamental for controlling virus replication or could be associated with persistent disease if ineffective (5, 6). However, the consequences of SARS-CoV-2 infection on the GI immune system and the local ability to respond to viral infection in severe disease is currently unknown.

The intestinal immune system is highly compartmentalised (7). Immune responses can be initiated in gut-associated lymphoid tissue (GALT) (8). Activated B and T cells generated in GALT acquire specific receptors, such as α4β7, CCR9 and CCR10 that allow them to home to lamina propria following circulation via lymphatics and the blood (9-11).

Peyer’s patches (PP) are clusters of GALT concentrated in the terminal ileum. A common feature of PP from early life in humans is the presence of germinal centre (GC) that are acquired in response to particulate antigens sampled from the gut lumen. The ensuing GC response generates lamina propria plasma cells secreting IgA that is transported into the gut lumen and that subsequently regulates the microbiota and maintains homeostasis (7, 12).

GC responses are regulated in part by transcription factor BCL6 (B-cell lymphoma 6) that is considered a marker for GC cells. It is known that GCs can be lost in lymph nodes and spleen in acute COVID-19, and this has been linked to diminishing of BCL6^+^ B and T cells in these tissues and blood (6). Whether GALT is similarly impacted is not known.

Here, the virus was quantified and localised in samples of gastrointestinal tract from patients who died with COVID-19 using reverse transcription quantitative PCR (RT-qPCR) and immunohistochemistry. By immunohistochemistry and imaging mass cytometry (IMC), the architecture and cellularity of PPs in the same samples were then explored in detail.

## Results

### Identification of SARS-CoV-2 in tissue samples along the GI tract

We first quantified and localised SARS-CoV-2 in formalin-fixed paraffin embedded (FFPE) samples of oesophagus, stomach, duodenum, ileum, colon, lungs and spleen from 7 males and 2 females who died after being diagnosed with COVID-19 (Supplementary Table 1). RT-qPCR analysis of N1 of SARS-CoV-2 nucleocapsid standardised to RNAse P revealed traces of the virus in most tissues from COVID-19 patients but not controls (Figure 1A-C). Immunohistochemistry with a cocktail of antibodies to the spike 2 glycoprotein and nucleocapsid of SARS-CoV-2 showed epithelial staining and punctate staining in subepithelial lamina propria. Double staining localised the punctate staining to CD68^+^ macrophages (Figure 1D). No virus staining was observed in lymphoid tissues (Figure 1D).

**Figure 1.**
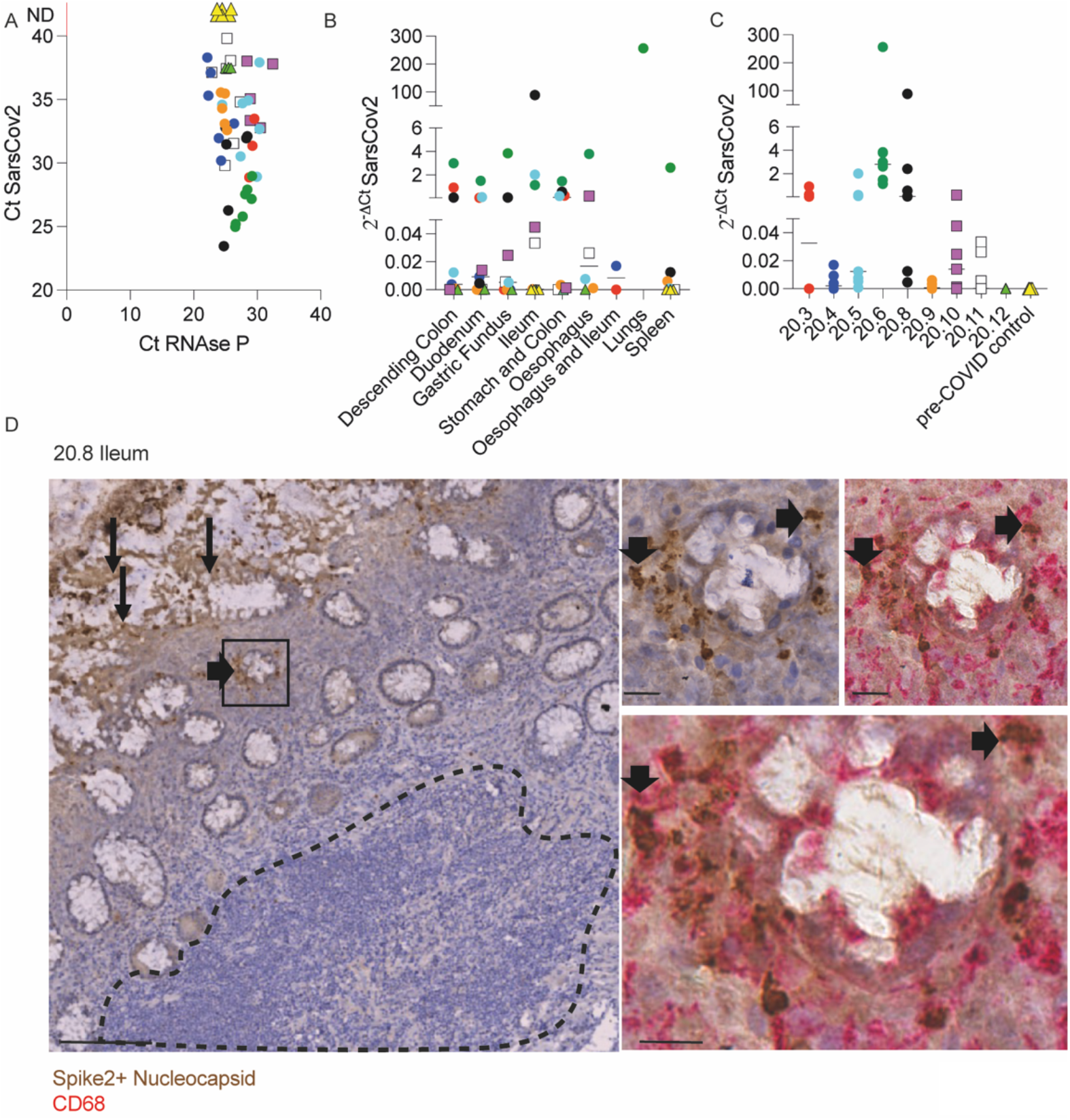
Identification of SARS-CoV-2 in tissue samples along the GI tract. (A-C) Evaluation of SARS-CoV-2 presence using RT-qPCR in different FFPET samples from 9 COVID-19 deceased patients and 4 pre-COVID patients used as controls. Dots represent the viral RNA level from each sample and were coloured by patient. The lines represent the median values *per* organ (B) or patient (C). (E) IHC for SARS-CoV-2 spike and nucleocapsid proteins in an ileal sample from patient 20.8 showing the virus presence in the epithelium (panel on the left) and sub-mucosal macrophages (panels on the right). Scale bar for image on the left: 100μm. Sale bars for images on the right: 20μm. The dashed line delimitates the GALT.

Therefore, in severe infection, SARS-CoV-2 is distributed along the digestive tract where it is localised mainly in epithelium and in subepithelial macrophages.

### Peyer’s patches from COVID-19 patients are depleted of germinal centres

In order to investigate better the architecture of PPs, ileal samples were initially double stained with anti-CD45RB that is expressed by T and the B cells on the periphery of lymphoid tissues, but not GC B cells (13) and anti-CD10 that stains the GC (14). The CD10:CD45RB ratio was significantly reduced in COVID-19 patients compared to controls, irrespective of the local levels of viral RNA measured by RT-qPCR (Figure 2A-B). Depletion of GC was therefore independent of the presence of local virus (Figure 2A).

**Figure 2.**
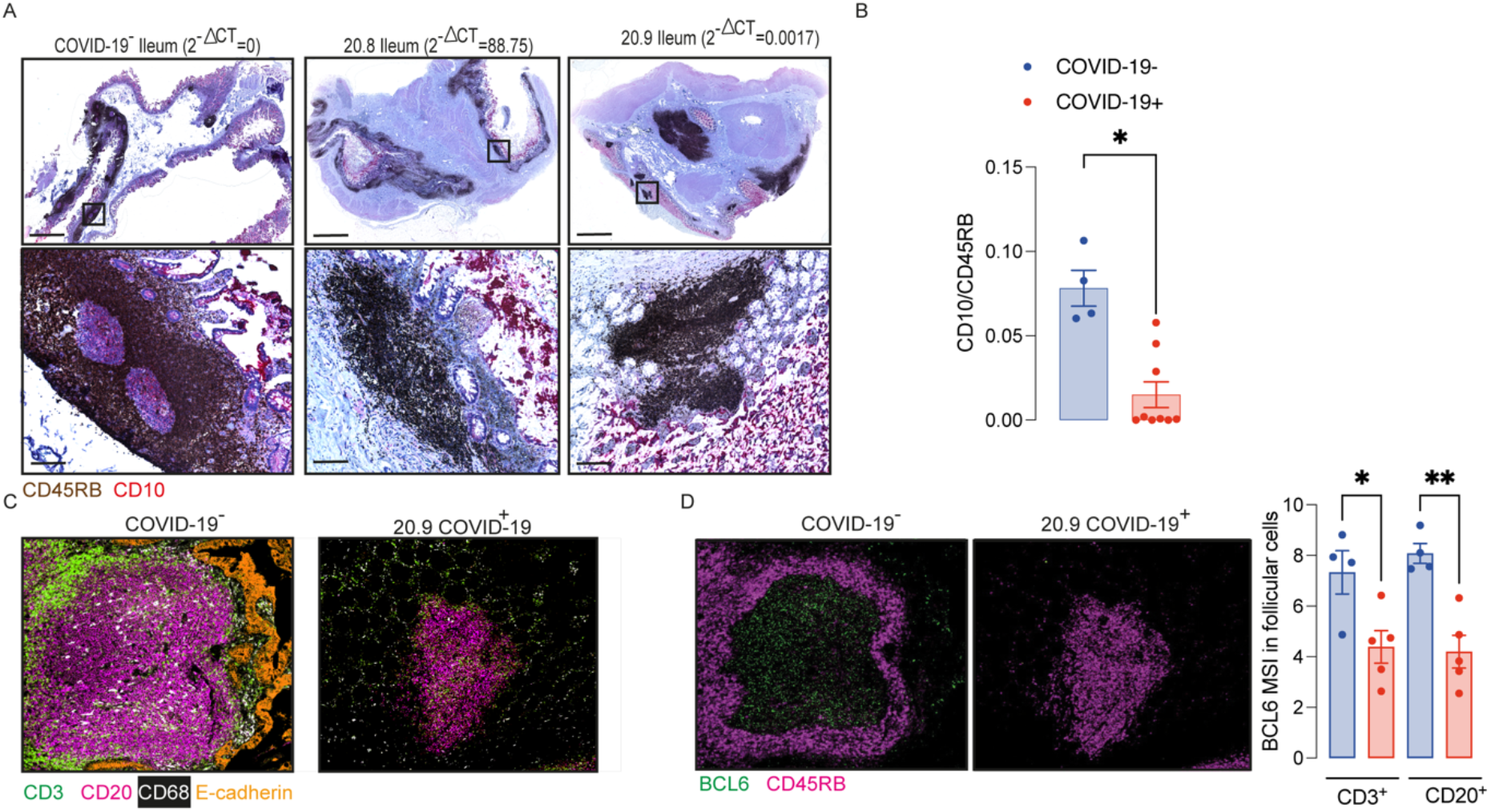
Peyer’s patches (PP) from COVID-19 patients lose germinal centre. (A-B) IHC for CD45RB (brown) and CD10 (red) in ileal FFPE samples from COVID-19 patients with different local levels of SARS-CoV-2 viral RNA. Images on the top show the whole sections and ileal follicles are highlighted in each of the bottom images. The images on the left represent a sample from one control; the images in the middle represent a sample from a COVID-19 patient with high local levels of viral RNA; and the images on the right represent a COVID-19 patient with low local levels of viral RNA. (B) CD10:CD45RB area ratio. Data is shown as mean± SEM. (n=4 controls and 9 patients). Two tailed Mann-Whitney t test. *P<0.05. (C) Representative images from histoCAT showing CD3 (green), CD20 (magenta), E-cadherin (orange) and CD68 (white) signals in ileal samples. (D) Representative images from histoCAT showing CD45RB (magenta) and BCL6 (green) signals in PP follicle on the left, and mean BCL6 signals in T and B cells on the right. Data is shown as mean± SEM. (n=4 controls and 5 patients). Kruskal-Wallis followed Dun’s post-test in D. *P<0.05.

IMC was used to characterize the cellularity of the ileal PP from 5 *post mortem* samples from COVID-19 patients and 4 controls including one from a *post mortem* and 3 surgical samples. Sections were stained with a cocktail of 22 antibodies (Supplementary Table 2) and areas of lymphoid tissue were ablated on a Hyperion imaging system (Fluidigm, South San Francisco, CA). The acquired raw images were visualized in histoCAT (15) and cells were then segmented using a pipeline based on pixel classification of multi-channel images using Ilastik (16) and Cell Profiler (17). The mean signal intensity (MSI) for each channel corresponding the antibody staining was extracted from individual cells and normalized between values of 0 and 100 before cell classification and heatmap validation (Supplementary Figure 1-3). Preliminary unsupervised clustering analyses by Seurat (18) and Phenograph(15) were not able to robustly identify the fundamental cell populations. Therefore, the cell classification was achieved using a basic gating strategy.

Cells were specifically selected from the lymphoid tissue in the PP and splenic white pulp for subsequent analysis (Supplementary Figure 4). IMC confirmed that the structure of the PP was disrupted in patients with COVID-19. Zonation of B cells and T cells was lost (Figure 2C). Expression of the GC-associated BCL6 transcription factor was reduced in T and B cells of follicular area from COVID-19 samples compared to those from controls (Figure 2D).

### Analysis of cellularity and cellular interactions in PP from COVID-19 patients

The relative numbers of T and B cells, CD4^+^ T cells, CD8^+^ T cells, CD4^+^FoxP3^+^ T cells and the mean signals for PD-1, CD27 and CD45RO in T cells were similar between COVID-19 patient and control samples (Supplementary Figure 5 and Figure 3A-D). The relative proportion of macrophages was higher in follicles of COVID-19 samples compared to controls (Figure 3E), although the percentages of CD14^+^CD16^-^ or CD14^+^CD16^+^ or CD14^lo^CD16^+^ macrophages were similar between groups (Figure 3F).

**Figure 3.**
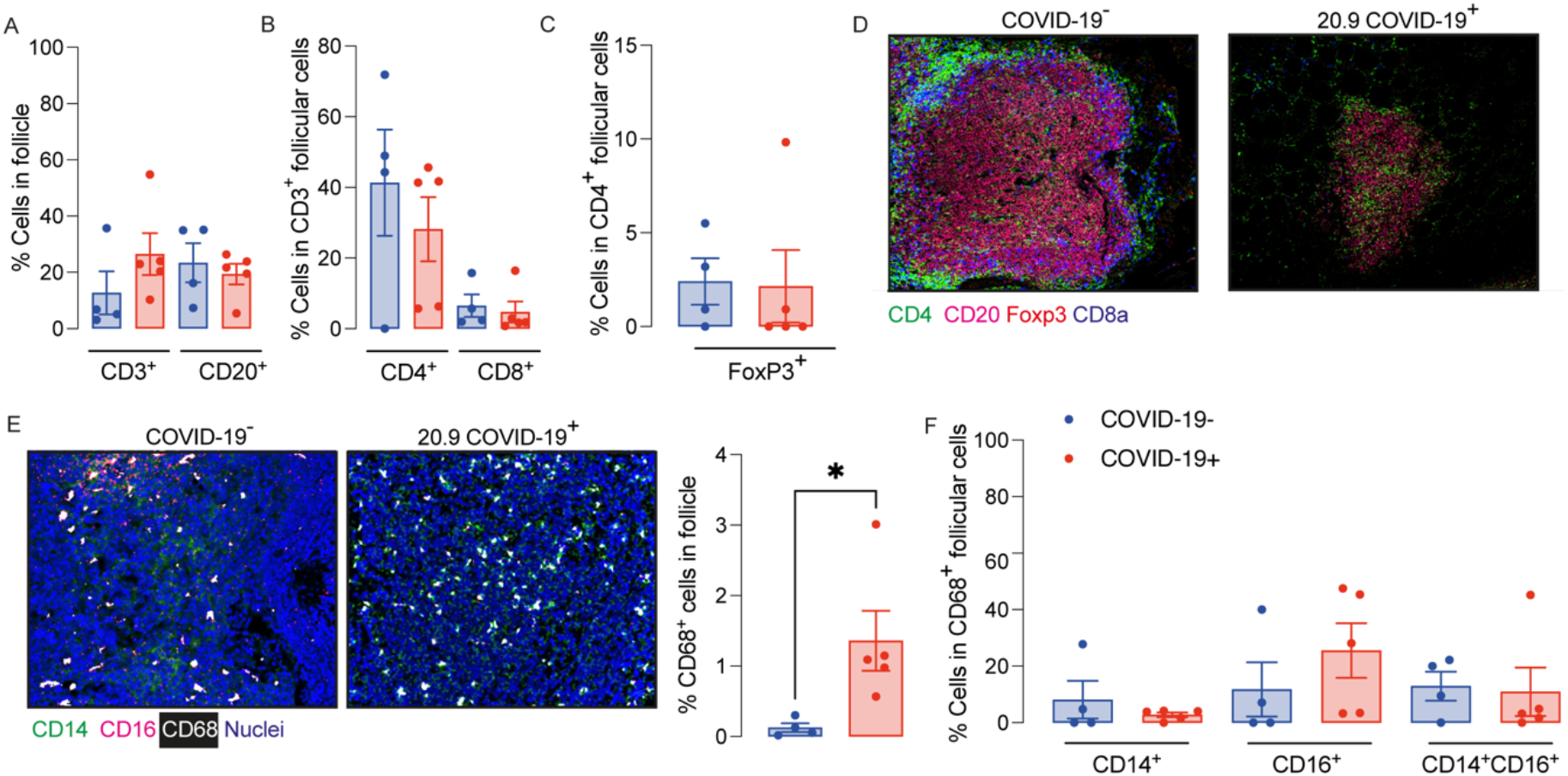
Enhanced relative numbers of macrophages in ileal follicles in Peyer’s patches (PP) of COVID-19 patients. (A-C) Percentages of follicular T and B cells, CD4^+^, CD8^+^ and CD4^+^FoxP3^+^ T cells in ileal Peyer’s Patch (PP) from COVID-19^-^ and COVID-19^+^ patients. (D) Representative images from histoCAT showing CD4 (green), CD8a (blue), FoxP3 (red) and CD20 (magenta) signals in PPs. (E) Representative images from histoCAT showing CD68 (white), CD14 (green) and CD16 (magenta) signals in PPs on the left, and mean CD68 signals on the right. (F) Percentages of follicular CD14^+^, CD16^+^ and CD14^+^CD16^+^ cells from CD68^+^ cell population. Data is shown as mean± SEM. (n=4 controls and 5 patients). Kruskal-Wallis followed Dun’s post-test in D and two tailed Mann-Whitney t test in E. *P<0.05.

The area of the ablated regions occupied by the lymphoid tissue was comparable between COVID-19 samples and controls (0.08±0.028 vs 0.09±0.021 mm^2^). However, the cellular density was significantly reduced in lymphoid tissue in COVID-19 (Figure 4A).

**Figure 4.**
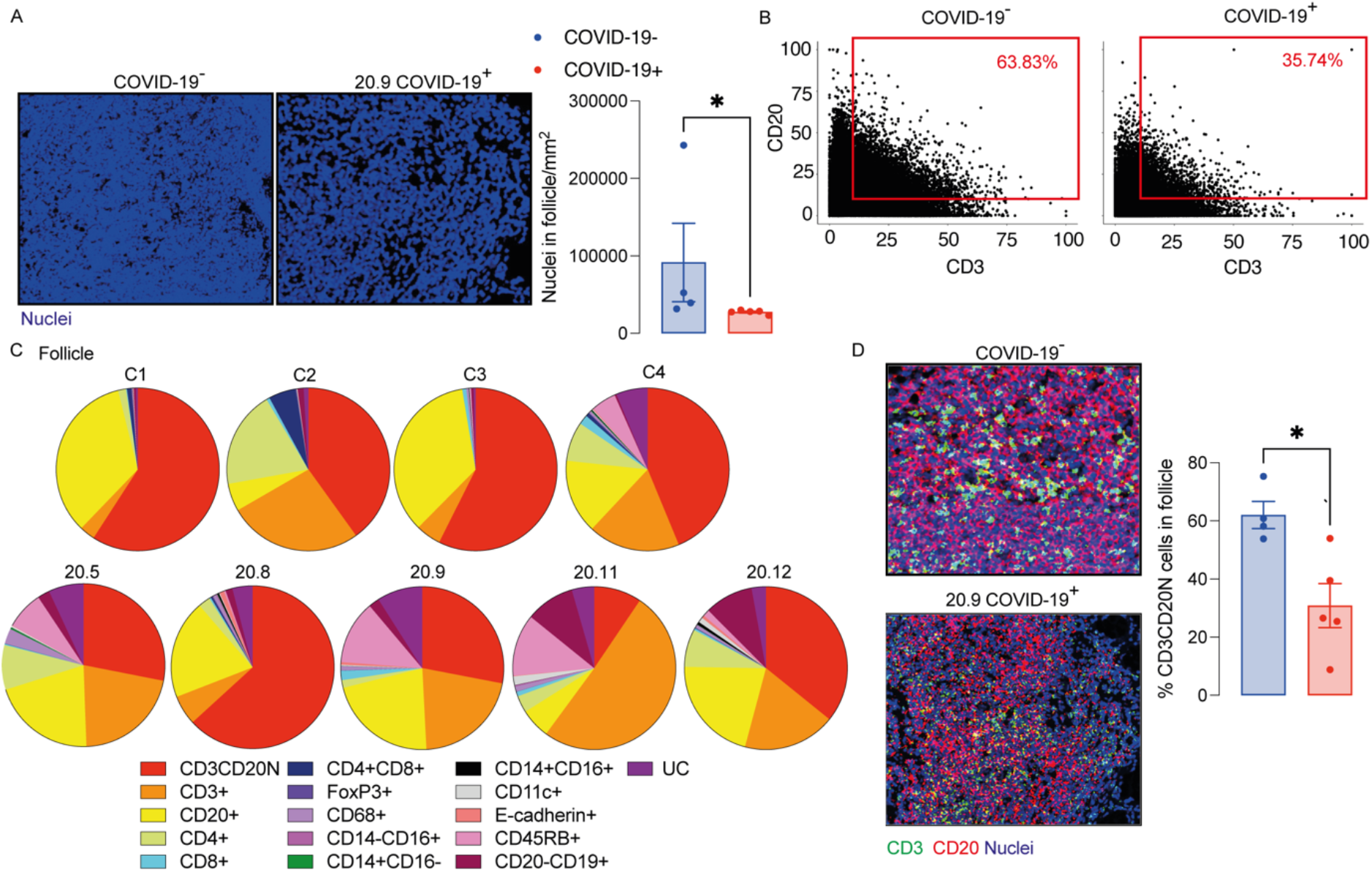
Decreased T and B cell interaction in ileal follicles in Peyer’s patches (PP) of COVID-19 patients. (A) Representative images from histoCAT showing nuclear density in ileal follicles in Peyer’s Patch (PP) from COVID-19^-^ and COVID-19^+^ patients on the left; and mean data on the right. (B) Dot-plots showing the interaction between T and B cells in ileal follicles. (C) Percentages of different cellular types in follicles from each control and COVID-19^+^ patient. CD3CD20N: T and B cell neighbours. UC: unclassified cells. (D) Representative images from histoCAT showing CD3 (green) and CD20 (red) merged signals (yellow) on the left the mean of proportions of CD3CD20 neighbours (CD3CD20N) on the right. Data is shown as mean± SEM. (n=4 controls and 5 patients). Mann-Whitney t test in A and D. *P<0.05.

As a surrogate measurement for T cell/B cell interaction, we identified segmented cells that gave membrane signal for both B cells and T cells and designated these cells CD3/CD20 neighbours (CD3CD20N). Proportionately fewer CD3CD20N were observed in the PP of COVID-19 samples compared to controls (Figure 4B-D). The lack of proximity of T cells and B cells in COVID-19 samples could also be observed by mixing of CD3 and CD20 signals in a single pixel in the follicle images (Figure 4D).

Considering the significant depletion of GC and reduced T cell/ B cell interaction in the ileal follicles in PP of COVID-19 patients, we next evaluated the CD27 and CD74 expression by B cells. The presence of CD27^+^CD20^+^ B cells and CD74^hi^CD20^+^ B cells were significantly reduced in PP from patients with COVID-19 (Figure 5A-B) compared to controls.

**Figure 5.**
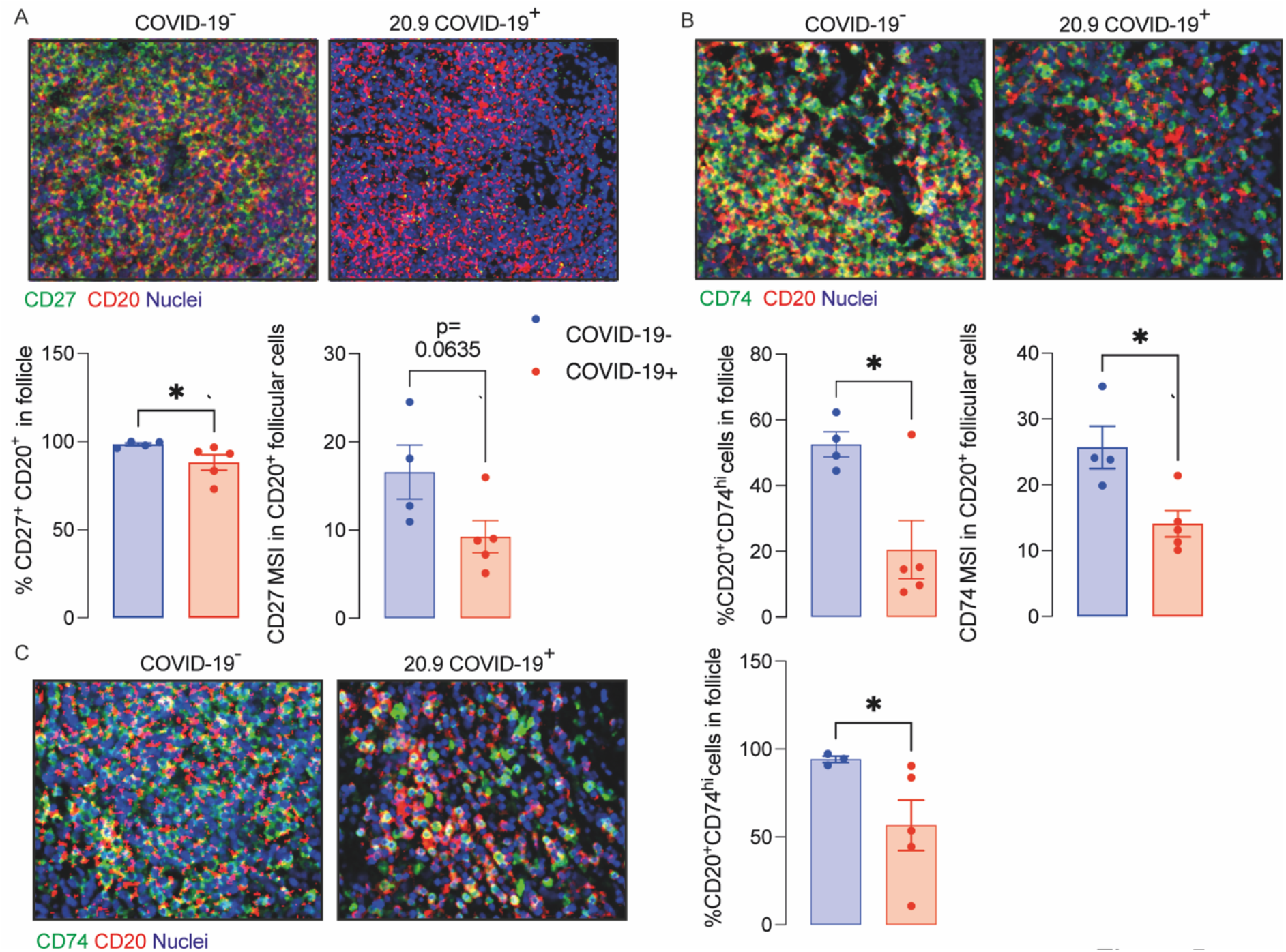
Decreased memory and antigen-presenting B cells in ileal follicles in Peyer’s patches (PP) of COVID-19 patients. (A) Representative images from histoCAT showing CD27 (green) and CD20 (red) in ileal follicles in Peyer’s Patch (PP) from COVID-19^-^ and COVID-19^+^ patients on the top. The percentage of CD27^+^CD20^+^ cells and mean signal of CD27 in B cells on the bottom. (B) Representative images from histoCAT showing CD74 (green) and CD20 (red) in ileal follicles on the top. The percentage of CD74^hi^CD20^+^ cells and mean signal of CD74 in B cells on the bottom. (C) Representative images from histoCAT showing CD74 (green) and CD20 (red) in white pulp from spleen on the left. The percentages of CD74^hi^CD20^+^ cells and mean signal of CD74 in B cells on the right.

**Figure 6.**
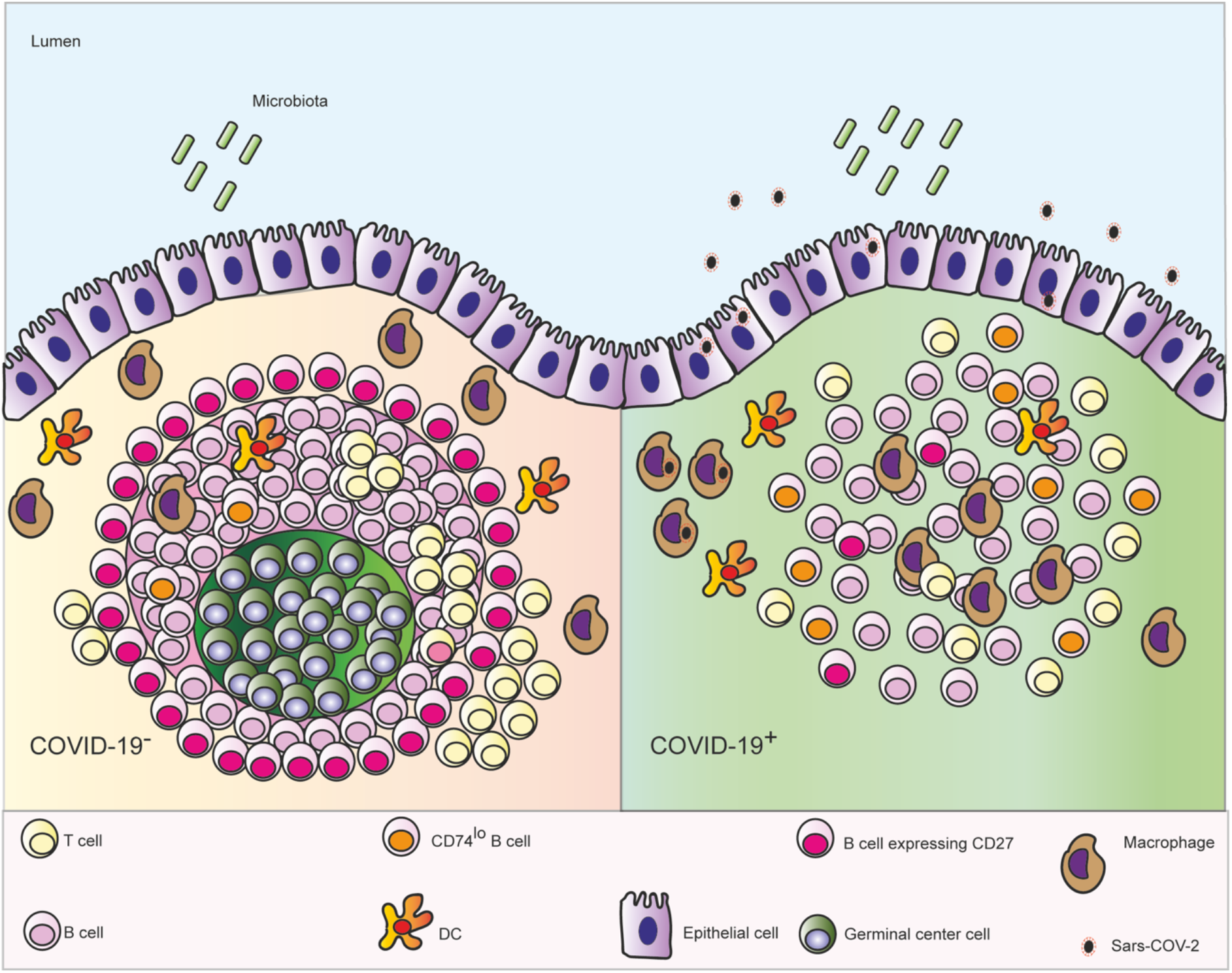
Schematic depicting the microanatomical features identified in ileal *post mortem* samples from patients who died with COVID-19: depletion of the germinal centre (GC) in the Peyer’s patches, enhanced numbers of follicular macrophages, decreased interaction between B and T cells, fewer CD27^+^ memory B cells and lower expression CD74 on B cells.

The data extracted from the lamina propria was highly variable between samples and therefore not described here.

Similar findings to those in PP, including the deficient CD74^hi^ B cells, were observed in splenic white pulp (Figure 5C), although the total populations of plasma cells (CD19^+^CD20^-^), T, B cells and memory T cells (CD45RO^+^) were comparable to controls (Supplementary Figure 6).

In summary, the structure and cellularity in PP of deceased COVID-19 patients is severely altered independent of the local levels of the virus. Such disturbed architecture could be related to changes in IgA production and microbiota described by previous studies(19, 20).

## Discussion

Infection with SARS-CoV-2 causes a range of symptoms including GI manifestations (3). Here we observed GI epithelial cells and subepithelial macrophages containing virus in patients who died with COVID-19. We also showed that microanatomy of PP of COVID-19 patients was severely affected by the disease, which was independent of the local levels of viral RNA.

We observed depletion of GC in the PP of patients who died with COVID-19. Depletion of GC in spleen and lymph nodes in *post mortem* samples from patients with COVID-19 has been reported previously (6). This was linked to failure of BCL6-expressing T follicular helper cells to support GC formation. In the present study a depletion of BCL6-expressing B and T cells in PP was also observed in COVID-19. Unlike in the spleen and lymph nodes that are commonly quiescent in the absence of infection, GC are constitutively present and clearly visible in PP from early stages of life (21). Our data therefore suggest that existing GC can become diminished in severe COVID-19. In addition, GC have reduced potential to form *de novo* through deficiencies in T cell B cell interactions.

A previous study has shown a decreased number of CD4^+^ T cells and CD19^+^ B cells expressing α4β7 integrin in blood of patients with severe COVID-19, even after remission of the symptoms (22). This integrin is imprinted in GALT and is essential for homing of cells generated in the PP back to the lamina propria (23). Lack of cells expressing α4β7 integrin in blood is consistent with compromised potential for immune induction and generation of effector cells in GALT observed here.

Changes in PPs were observed even when only trace levels of local virus were detected in the tissues. It therefore seems likely that changes observed in PP are related to the systemic inflammation rather than the virus *per se*. Lymphopenia observed in these patients and in other studies (24) supports this concept. Indeed, a study in mice showed that LPS injection resulted in a significant reduction in numbers of T and B lymphocytes in PP, which was attributed to an increased apoptosis rate (25). Another group showed thermal injury-plus-sepsis or sepsis alone in rats lead to a suppressed CD4^+^ T proliferation/IL-2 production and a substantial down-modulation of lymphocyte survival in mesenteric lymph nodes (26).

B cells expressing CD27 were depleted in GALT of COVID-19 patients evaluated here. This is consistent with depletion of B cells with the phenotype CD27^+^IgM^+^IgD^+^ seen in blood (24, 27). This is also consistent with a depletion of CD27^+^ B cells in blood in sepsis (28). B cells in PP of COVID-19 patients expressed lower levels of CD74 than in controls. Lower antigen-presentation capacity previously reported in patients with sepsis with reduced HLA-DR expression in monocytes inversely correlated with the severity of multi-organ damage (29).

The decreased cell density of the PP and depletion of the GC in ileal follicles of patients with COVID-19 is consistent with impaired T and B cell interaction, which could contribute to failure to generate a long-term response to local antigens and contribute to dysbiosis (19, 30).

In conclusion, patients with severe COVID-19 show significant impaired architecture and cellularity of PP. The resulting poor local immunity could contribute to dysbiosis. Our findings also suggest that oral vaccination to prevent COVID-19 disease could not be effective if patients were already ill, since the gut immune system is compromised with features indicating that they would lack the ability to mount an efficient immune response.

## Material and Methods

### COVID-19 patients

Formalin fixed paraffin embedded (FFPE) samples including samples of oesophagus, stomach, duodenum, ileum, colon, lung and spleen were obtained from 9 patients who died with severe COVID-19. Sex, age, body mass index (BMI), symptoms at admission, time to death and some laboratorial parameters are described in Supplementary Table 1.

### Pre-pandemic negative samples

A *post-mortem* ileal sample obtained from a patient with metastatic lung adenocarcinoma and 3 surgical resections of ileum and spleen obtained from anonymous donor’s pre-pandemic were used as COVID-19^-^ controls.

### RT- qPCR

Total RNA was extracted from 10μm-sections of FFPE tissues using a commercial kit (High pure FFPET RNA isolation kit, Roche, Cat.6650775001) and resuspended in 25μl final volume. RT-qPCR reactions were performed with Taq-Man™ Fast virus 1-step master mix (ThermoFisher Scientific, Cat. 4444436) and primer-probe sets targeting SARS-CoV-2 nucleocapsid (N1 set) and human RNAse P (CDC 2019-Novel Coronavirus (2019-nCoV) RT-PCR diagnostic panel (Centers for Disease Control and Prevention, Cat. 2019-nCoVEUA-01).

### Immunohistochemistry (IHC)

5μm- sections of FFPE tissues on SuperFrost Plus adhesion slides (ThermoScientific, Cat. 10149870) were deparaffinized using 3 solutions of absolute m-xylene or Histoclear II histology (SLS, Cat. NAT1334), rehydrate in 90%, 80%, 70% ethanol solutions and DPBS (ThermoFisher, Cat.14190169). After then, the slides were immersed in an antigen retrieval solution pH 9.0 (Agilent Dako, S236784-2) and kept in a pressure cooker for 2 minutes. The cells into tissues were permeabilized in a 0.1%Tween20 solution for 5 minutes. The blocking, staining with primary and secondary antibodies were realized according to the manufactures of the following IHC kits: Mouse and rabbit specific HRP/DAB IHC detection Kit micro-polymer (Abcam, Cat.ab236466) and ImmPRESS[R] Duet double staining kit anti-rabbit AP/anti-mouse HRP (Vector laboratories, Cat.mp7724). Anti-SARS-CoV-2 spike antibody, targeting the S2 subunit, and anti-SARS-CoV-2 nucleocapsid (1:300) antibody were acquired from Insight Biotechnology (Cat. GTX632604) and BioserverUK (Cat.BSV- COV-AB-13), respectively. Anti-CD68 was obtained from Cell signalling (Cat. 7643T). All samples were stained with haematoxylin.

### Antibodies

Antibodies conjugated with metals were acquired from Fluidigm and the catalogue numbers are displayed in Supplementary Table 2. Anti-CD11c, anti-CD45RB, anti-CD103, and anti-ACE2 were labelled in house with metals using commercial kits (MaxPar labelling Kits, Fluidigm) according to the manufacturer’s instructions. All the antibodies used were validated by IHC.

### Imaging mass cytometry

Deparaffinization, rehydration, antigen retrieval and permeabilization were the same as described previously. After permeabilization, unspecific epitopes were blocked in a solution made of 10%BSA, 0.1%Tween20, 1:20 Human TryStain FcX (Biolegend, Cat. 422301) and 1:20 Kiovig (5mg/ml solution, Baxter). The samples were incubated in a wet chamber with the antibody cocktail overnight and washed in DPBS plus Tween 0.1%. The nuclei were stained with a 1μM solution of Cell-ID™ Intercalator-Ir (Fluidigm, Cat.201192B) in DPBS. Finally, the samples were desalinized in milli-Q water and were kept in a dry place protected from any sources of oxidation until the tissue ablation. The tissues were visualized under a light microscope and after choosing a region of interest (where the lymphoid tissue was present), data was acquired on a Hyperion imaging system coupled to a Helios Mass Cytometer (Fluidigm) tuned with 3 element tuning slide, at a laser frequency mean of 200Hz and 6dB power.

### Imaging mass cytometry analyses

The acquired images were visualized in histoCAT(15) and segmented using Bodenmiller’s group pipeline (https://doi.org/10.5281/zenedo.3841961). The data was normalized in GraphPad Prism software v9.0 (GraphPad Software, Inc., La Jolla, CA) and cell populations defined in R studio according to the MSI parameters (Supplementary Figures 3-5). The x and y nuclear localization coordinates of all cells were plotted and the follicle cells or white pulp cells were selected using a custom R script to manually draw around the follicle or white pulp following the B cells signal visualized in histoCAT (Supplementary Figure 6). The % of area occupied by the follicle was determined in Image J (version 1.0 for Mac OS X) using the image selected in R studio. The follicular area was converted in mm^2^ by taking the size of the total ablated area and the % of area occupied by the follicle.

### Statistics

Analyses were performed using GraphPad Prism software v9.0 (GraphPad Software, Inc., La Jolla, CA). Data are reported as mean ± SEM. Comparisons were undertaken using Kruskal-Wallis followed by Dunn’s post-test or Mann Whitney t-test. *P<*0.05 was considered significant.

## Author Contributions

J.S.supervised the study. J.S. S.N. and S.C.T. conceived the study and contributed to experimental design. S.C.T., S.P., L.M., J.G.M., K.T., C.B. performed the experiments. S.C.T., F.S., N.P. and M.P. analysed the data. J.S. and S.C.T. interpreted data and wrote the manuscript. S.N, A.G. and M.S.H. provided critical intellectual input.

## Funding

This study was supported by UK Research and Innovation (UKRI) and the National Institute of Health Research (NIHR) via the UK Coronavirus Immunology Consortium. This research was also funded/supported by the National Institute for Health Research (NIHR) Biomedical Research Centre based at Guy’s and St Thomas’ NHS Foundation Trust and King’s College London and/or the NIHR Clinical Research Facility. The views expressed are those of the author(s) and not necessarily those of the NHS, the NIHR or the Department of Health and Social Care.

Human samples used in this research project were obtained from the Imperial College Healthcare Tissue Bank (ICHTB). ICHTB is supported by the National Institute for Health Research (NIHR) Biomedical Research Centre based at Imperial College Healthcare NHS Trust and Imperial College London. ICHTB is approved by Wales REC3 to release human material for research (17/WA/0161), and the samples for this project (give approved project number R20044) were issued from sub-collection reference number MED_MO_20_011: Covid 19 Post Mortem Research.

## Acknowledgements

We thank Professor Mathew Brown for his intellectual input and for allowing us to use his laboratory facilities. We also acknowledge Kishan Joshi for supporting the acquisition of samples studied.

## Notes

### Competing Interest Statement

The authors have declared no competing interest.

