## Supplementary material for "Disrupted Peyer’s patch microanatomy in COVID-19 including germinal centre atrophy independent of local virus"

### Supplementary figures and tables

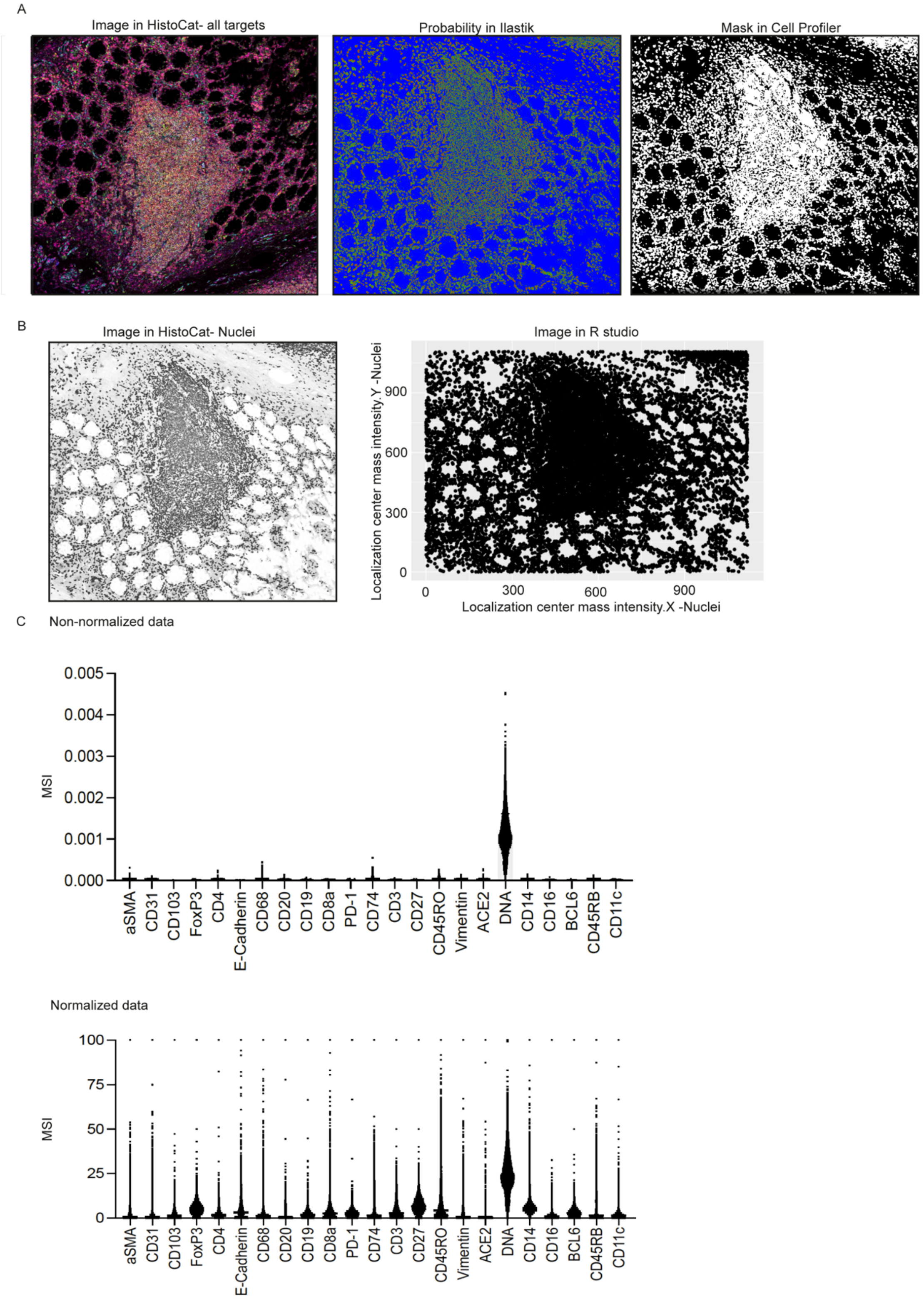


**Supplementary Figure 1. Cell segmentation and data normalization.** Ileal FFPE samples analysed by imaging mass cytometry. (A) The representative image on the left shows an ileal follicle visualized in histoCAT containing the signal from all studied channels. The image on the middle shows the probabilities obtained in Ilastik for the nuclei (red), membranes (green) and background (blue). The image on the right shows the cell mask (nuclei+ membrane in white) of all identified cells (objects ranging from 5 to 21μm). (B) HistoCAT image of the nuclear signal is shown on the left. On the right is represented the same image reconstructed in R studio using the x-y nuclear localization coordinates. (C) The image on the top shows the mean of signal intensity (MSI) for cells in each channel for the patient 20.9 before normalization. The normalized data for the same patient is shown on the bottom.

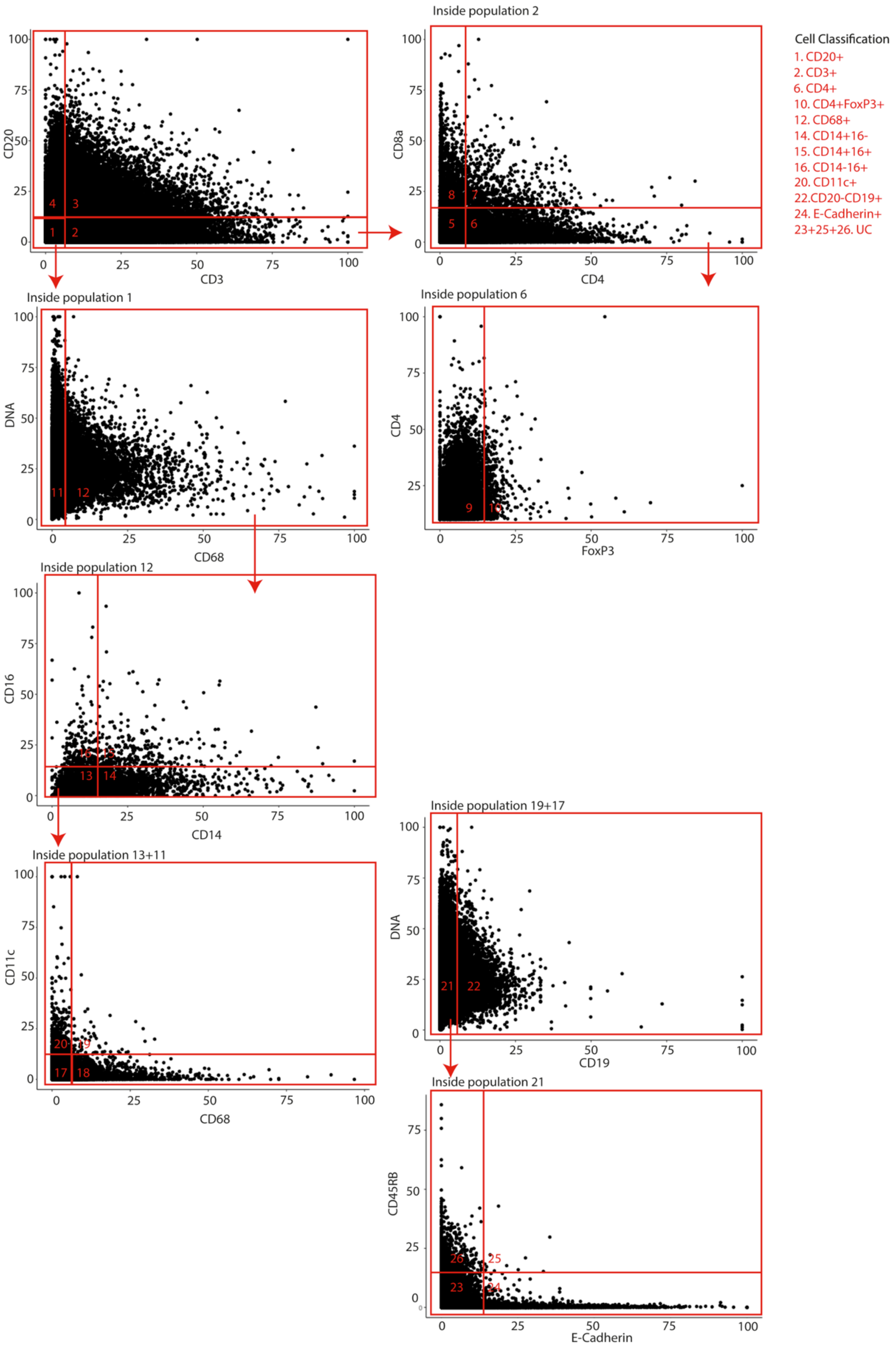


**Supplementary Figure 2. Gating strategy used for cell classification.** Ileal and splenic FFPE samples analysed by imaging mass cytometry. Cells were plotted according to signals for CD3

and CD20 and 4 populations primarily obtained (1. $CD3^-CD20^-$; 2. $CD3^+$; 3. CD3CD20N;4. $CD3^-CD20^+$). Inside the population 2, $CD4^+$, $CD4^+CD8^+$ and $CD4^+$ T cells were classified. The $CD4^+$ T cells were further classified in $CD4^+FoxP3^+$ cells considering the signal for FoxP3. Inside the population 1, $CD68^+$ cells were classified and further divided in $CD14^+CD16^-$, $CD14^-CD16^+$ and $CD14^+$ $CD16^+$ cells. The $CD14^-CD16^-$ cells and $CD68^-$ cells were joined in a new file and plotted according to signals for CD11c and CD68 to obtain $CD11c^+$ cells. The populations $CD11c^+CD68^+$ and $CD11c^-CD68^-$ were joined in a new file and plotted according to signals for CD19 to obtain $CD20^-CD19^+$ cells. The $CD19^-$ cells were plotted according to signals for CD45RB and E-cadherin to obtain $CD45RB^+$ and E-cadherin$^+$ cells. All the remaining cells were considered unclassified.

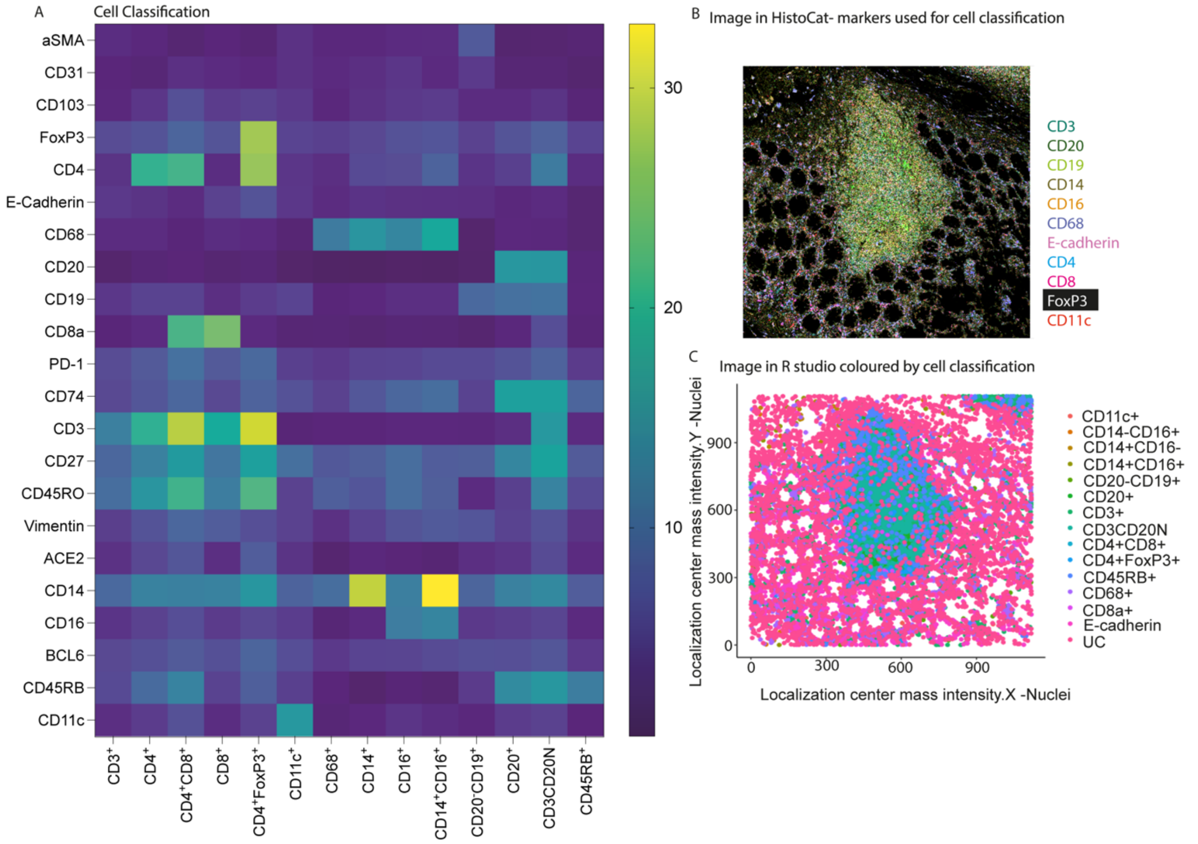


**Supplementary Figure 3. Validation of the cell classification.** Ileal FFPE samples analysed by imaging mass cytometry. (A) The mean of intensity signal for each channel was represented in a Heatmap according to the cell classification obtained as shown in Supplementary Figure 2. (B) HistoCAT image containing the signals for CD3 (teal), CD20 (fern), CD11c (red), CD14 (asparagus), CD16 (orange), CD19 (light green), CD68 (orchid), CD4 (light blue), CD8 (magenta), FoxP3 (white), CD45Rb (salmon) and E-cadherin (pink). (C) The same image shown in B was reconstructed in R studio according to the x-y nuclear localization coordinates, and the cells labelled according to the cell classification obtained in giving Supplementary Figure 2.

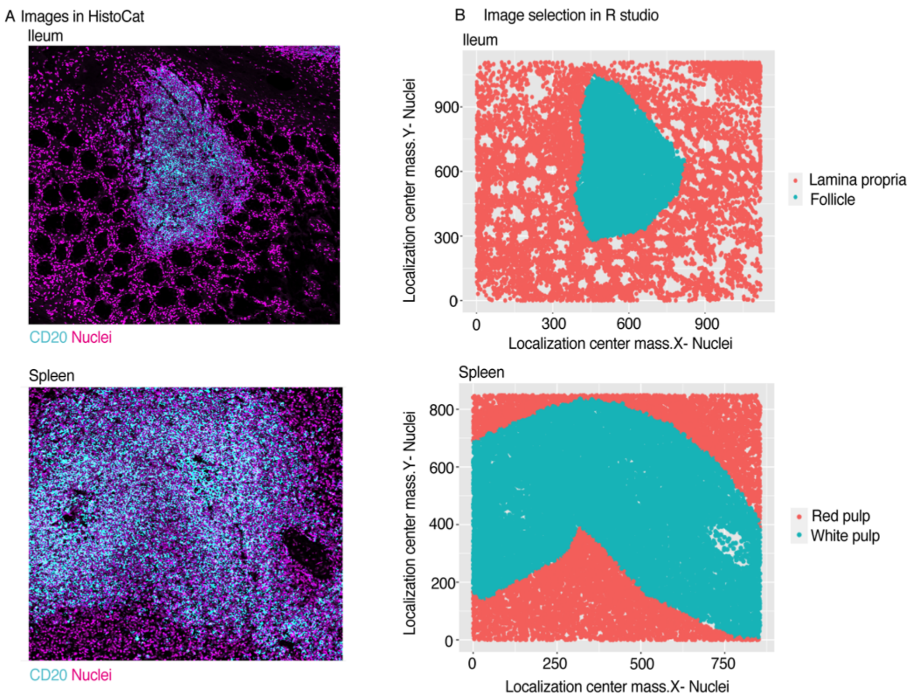


**Supplementary Figure 4. Selection of follicular and white pulp areas in ileum and splenic samples respectively.** (A) The representative image on the top shows an image of an ileum sample from the patient 20.9 visualized in histoCAT containing the signal for nuclei (magenta) and CD20 (light blue). The representative image on the bottom shows an image of a splenic sample from the same patient. (B) Representative images in A were reconstructed in R studio using the x-y nuclear localization coordinates and follicular and white pulp areas (green) manually defined based on the CD20 signal.

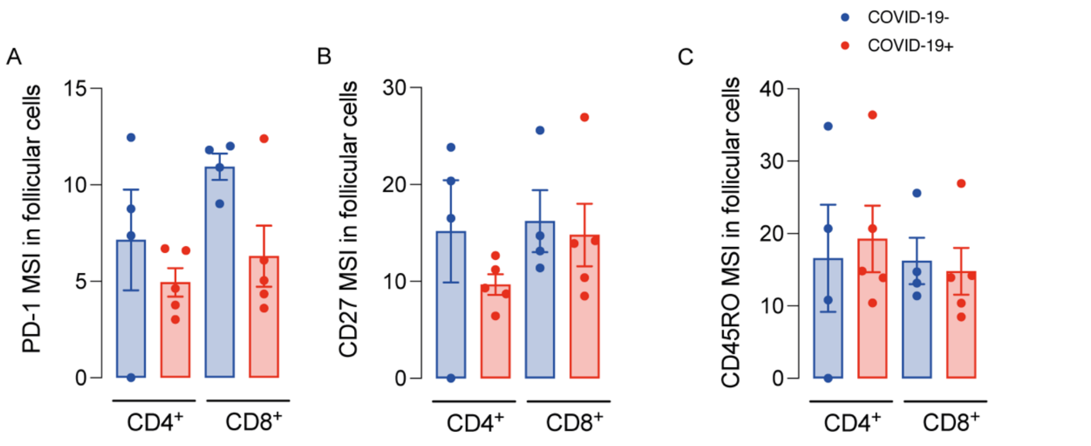


**Supplementary Figure 5. Signals for PD-1, CD27 and CD45RO in ileal follicular T cells are comparable between COVID-19⁻ and COVID-19⁺ patients.** FFPET samples analysed by imaging mass cytometry. (A-C) Mean of signal intensity for PD-1 (A), CD27(B) and CD45RO(C) in CD4⁺ and CD8⁺ follicular T cells in ileum.

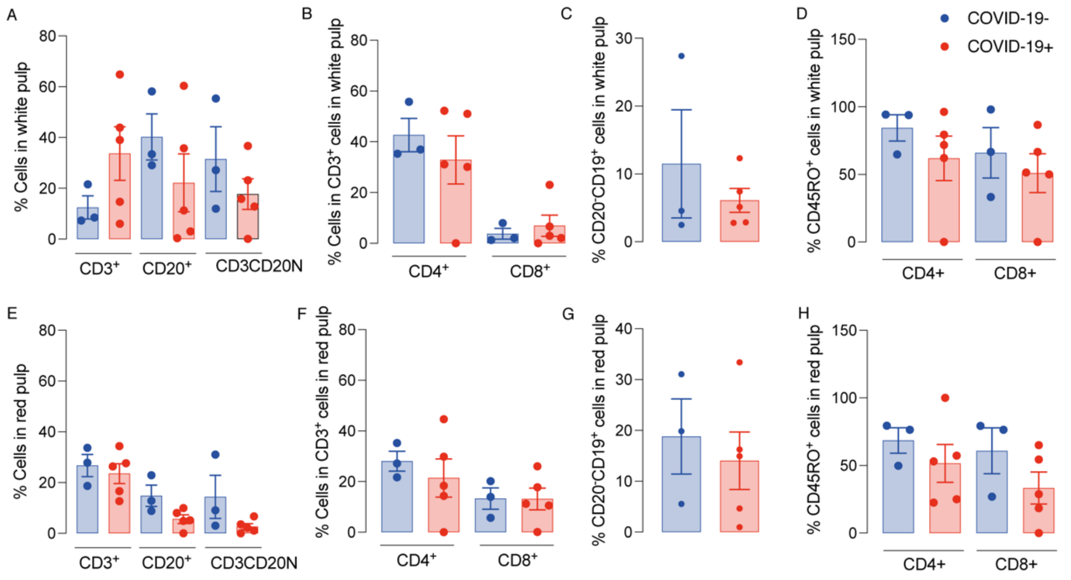


**Supplementary Figure 6. Relative numbers of splenic T and B cells are comparable between COVID-19⁻ and COVID-19⁺ patients.** Relative numbers of T cells (CD3⁺), B cells (CD20⁺), plasma cells (CD20⁻CD19⁺), memory CD4⁺ and CD8⁺ T cells (CD45RO⁺) in white pulp (A-D) and red pulp (E-H) of splenic FFPE samples analysed by imaging mass cytometry.

| Patient ID (gender) | Body Mass Index (kg/m²) | Age (years) | Total lymphocytes (x10⁹/L) | CRP (mg/L) | Hemoglobin (g/L) | Time symptoms to death (days) | Symptoms at admission |
|---|---|---|---|---|---|---|---|
| 20.3 (M) | 31 | 79 | 0.4 | 352.2 | 94 | 23 | Fever, low $O_2$ saturation |
| 20.4 (M) | 18.3 | 97 | 0.6 | 143 | 135 | 23 | General malaise |
| 20.5 (M) | 33.1 | 61 | | | | 10 | Cyanosis, acute chest pain |
| 20.6 (M) | 25.2 | 24 | 0.6 | 13.1 | 88 | 8 | Cough, fever, short breath |
| 20.8 (F) | 19.72 | 79 | 0.7 | 220 | 87 | 8 | Vomiting, diarrhea |
| 20.9 (M) | 35.86 | 64 | 0.7 | 416.9 | 119 | 13 | Low $O_2$ saturation |
| 20.10 (F) | 44.1 | 69 | 0.6 | 118.8 | 143 | 8 | Cough, fever, short breath |
| 20.11 (M) | 24.7 | 78 | 1.5 | | 135 | 12 | Cough, fever, short breath |
| 20.12 (M) | 48.8 | 22 | | 395 | 78 | 27 | Pneumonia, cerebral artery infarct |
| Control | 18.5-24.9 | - | 1.5-4.5 | 0-5 | 130-180 | NA | NA |
| Control PM (F) | - | 57 | - | - | - | - | Metastatic lung adenocarcinoma |

**Supplementary Table 1. Summary of patient's data.** Normality values are expressed on the control row. M: male, F: female, NA: non-applicable, CRP: C-reactive protein. PM: *post mortem*.

| Metal | Target | Dilution | Company (Cat. number) |
|---|---|---|---|
| 141Pr | alpha-SMA | 1:4000 | Fluidigm (3141017D) |
| 142Nd | CD19 | 1:100 | Fluidigm (3142014D) |
| 143Nd | Vimentin | 1:500 | Fluidigm (3143027D) |
| 144Nd | CD14 | 1:100 | Fluidigm (314025D) |
| 146Nd | CD16 | 1:200 | Fluidigm (3146020D) |
| 147Sm | BCL6 | 1:50 | Fluidigm (3147020D) |
| 148Nd | CD45RB | 1:1000 | Biolegend (310202) |
| 150Nd | CD11c | 1:300 | Abcam (ab216655) |
| 151Eu | CD31 | 1:300 | Fluidigm (3151025D) |
| 152Sm | CD103 | 1:100 | Abcam (ab254201) |
| 155Gd | FoxP3 | 1:300 | Fluidigm (3155016D) |
| 156Gd | CD4 | 1:200 | Fluidigm (3156033D) |
| 158Gd | E-cadherin | 1:3000 | Fluidigm (3158029D) |
| 159Tb | CD68 | 1:400 | Fluidigm (3159035D) |
| 161Dy | CD20 | 1:300 | Fluidigm (3161029D) |
| 162Dy | CD8a | 1:800 | Fluidigm (3162034D) |
| 165Ho | PD-1 | 1:50 | Fluidigm (3165039D) |
| 166Er | CD74 | 1:100 | Fluidigm (3166018D) |
| 170Er | CD3 | 1:300 | Fluidigm (3170019D) |
| 171Yb | CD27 | 1:300 | Fluidigm (3171024D) |
| 173Yb | CD45RO | 1:500 | Fluidigm (3173016D) |
| 176Yb | ACE2 | 1:100 | R&D(171608) |

**Supplementary Table 2. Antibodies used in Imaging Mass Cytometry (IMC).** Pr: praseodymium, Nd:neodymium, Sm: samarium, Eu: europium, Gd: gadolinium, Tb: terbium, Dy: dysprosium, Ho: holmium, Er: erbium, Yb: yttrium. CD: cluster of differentiation.
